# TRIM28 regulates pre-mRNA splicing via phosphorylation and SUMOylation networks

**DOI:** 10.1101/2025.07.07.663440

**Authors:** Vigor Matkovic, Cristian Prieto-Garcia, Thorsten Mosler, Kristina Wagner, Paul Hotz, Felix Haidle, Joost Schimmel, Nicole Blümel, Wojciech P. Galej, Alfred C.O. Vertegaal, Michaela Müller-McNicoll, Kathi Zarnack, Stefan Müller, Ivan Dikic

## Abstract

Pre-mRNA splicing is a highly regulated process orchestrated by splicing factors, cis-acting elements, and interconnected cellular processes such as transcription and chromatin remodeling. Here, we identify a novel regulatory axis involving phosphorylation and SUMOylation that governs the function of Tripartite motif-containing 28 protein (TRIM28) and its role in pre-mRNA splicing. We demonstrate that TRIM28 interacts with the spliceosomal protein USP39 in a phosphorylation-dependent manner, with non-phosphorylated TRIM28 promoting USP39 SUMOylation at defined lysine residues. This post-translational modification enhances USP39’s role within the U4/U6.U5 tri-snRNP complex. Functionally, TRIM28 knockdown induces widespread alterations in alternative splicing patterns, underscoring its importance in splicing regulation. Together, our findings uncover a mechanistic link between TRIM28-mediated post-translational modifications and the modulation of spliceosomal activity, offering new insights into how splicing decisions are integrated with cellular signaling pathways.

## INTRODUCTION

The regulation of cellular functions often relies on versatile proteins capable of orchestrating different biological pathways and influencing numerous aspects of cellular activity^1^. TRIM28 is one such protein, recognized for its extensive role in modulating essential processes through diverse interactions and post-translational modifications. Known also as TIF1β (transcriptional intermediary factor 1β) or KAP1 (KRAB-associated protein 1), TRIM28 acts as a polyvalent scaffold protein in different subcellular complexes regulating diverse cellular processes, including epigenetic modification and cell cycle control^2–11^. In epigenetics, TRIM28 is involved in transcriptional repression, forming complexes with KRAB domain-containing zinc-finger proteins. This interaction recruits chromatin-modifying enzymes, such as histone deacetylases and methyltransferases, to silence target genes^2,4,10^. Additionally, TRIM28 acts as a sensor of DNA damage responding through dynamic post-translational modifications. ATM kinase-driven phosphorylation of TRIM28 through Chk2 kinase on serine residues initiates a cascade of signaling events that promote DNA repair, cell cycle arrest, and apoptosis - crucial responses to maintain genomic integrity under genotoxic stress^4^. TRIM28 dissociates from heterochromatin-associated factors leading to relaxation of chromatin and activation of genes involved in DNA damage repair^12^. Research papers also suggest that TRIM28 may have intrinsic E3 SUMO ligase activity or at least, facilitate and mediate SUMOylation of target proteins^7^, which influences their (complex) stability, localization, and function. Moreover, auto-SUMOylation^13^ of TRIM28 that happens within its bromodomain, enhances its interaction with other co-repressor proteins and chromatin modifiers and only in the SUMOylated state, it can recruit the silencing complexes. Through SUMOylation, TRIM28 extends its regulatory influence across different cellular pathways^7^ and specificity and regulation can be achieved through interaction with specific substrates, domain-specific functions, regulation by post-translational modifications or context dependent activities^14–17^.

SUMOylation^18–20^ is a critical post-translational modification that involves the covalent conjugation of small ubiquitin-like modifier (SUMO) proteins to lysine residues on target proteins, thereby altering activity, localization or stability of cellular complexes^19,21,22^. This modification plays an essential role in regulating various molecular and cellular processes, including pre-mRNA splicing, fundamental step in gene expression. Splicing^23–31^ is catalyzed by the spliceosome, a large multi-subcomplex composed of small nuclear ribonucleoproteins (snRNPs) and a variety of splicing factors, which removes intronic regions from the precursors of messenger RNA transcripts and ligates exons to produce mature mRNA for translation into proteins. Dynamic assembly, activity and fidelity of the spliceosome are tightly regulated by different post-translational modifications that include phosphorylation^32–34^, ubiquitination^34^ and the SUMOylation.^35^ Through SUMOylation of specific spliceosomal proteins and regulatory factors, this modification can influence on splice site selection, the efficiency of splicing and alternative splicing decisions.^36,37^

The full impact of SUMOylation of spliceosomal proteins on splicing specificity and regulation remains still incompletely understood. SUMO E3 ligases like PIAS1 have been identified as involved in conjugation of SUMO to target proteins.^38^ Moreover, SUMO-specific protases (SENPs) contribute to the dynamics of SUMOylation by reversing this modification.^19^ Despite all these insights, the precise factors that facilitate the specificity of spliceosomal SUMOylation remain still unknown.

One of the important functional components of the spliceosome is USP39^39,40^, a protein involved in the maturation of the U4/U6.U5 tri-snRNP complex^41^, that is required for the formation of an active spliceosome and further splicing catalysis. Unlike typical deubiquitinating enzymes (DUB) ^42–44^, USP39 lacks catalytic residues and activity, yet it is indispensable for spliceosome assembly and function ^45–52^. USP39 has been implicated in regulating alternative splicing events, particularly for genes involved in cell division and signaling. It is involved in the splicing of mRNAs encoding AURKB (Aurora B kinase)^53^, an essential factor for mitotic spindle assembly and chromosome segregation. Depletion of USP39 disrupts spliceosome formation, resulting in defective splicing and subsequent mRNA degradation^49,54,55^, underscoring its significance in maintaining splicing fidelity^45,54^.

Here, we identify among all SUMOylated proteins, USP39 as highly SUMOylated spliceosomal protein and TRIM28 as a key interactor of USP39 that is controlling the SUMOylation status of USP39. Moreover, using minigene experiments we propose an influence of USP39 SUMOylation on the kinetics of splicing processes where SUMOylated USP39 causes conditions responsible for splicing stimulation.

Our results reveal a phosphorylation-dependent interaction between USP39 and TRIM28, with TRIM28 binding to USP39 occurring only in its non-phosphorylated state, while phosphorylation abrogates that interact and changes functional engagement of TRIM28. Altogether, our findings propose a new regulatory axis where TRIM28 modulates splicing dynamics through SUMOylation of USP39, linking post-translational modifications to spliceosome function and gene expression regulation.

## RESULTS

### Spliceosomal proteins are heavily SUMOylated and interact with USP39

To gain a deeper understanding of SUMOylation as a post-translational modification of splicing factors, we performed a series of mass spectrometry experiments to identify highly SUMOylated proteins within the global human proteome (Figure 1A). Notably, among the SUMOylated proteins identified, splicing factors stood out as highly SUMOylated (Figure 1A). GO term analysis of the proteins identified in Figure 1A confirmed that spliceosomal complexes are the among the highly enriched proteins, emphasizing the importance of SUMOylation in spliceosomal protein function (Figure 1B). Similarly, analysis of the biological processes associated with these SUMOylated proteins across the global proteome found RNA splicing and related gene ontology (GO) terms as highly enriched categories (Figure 1C). Among all hits, USP39 emerged as a highly SUMOylated spliceosomal protein (Supplementary Table S1). USP39 is an essential component of the U4/U6-U5 tri-snRNP splicing complex^56^.

**Figure 1.**
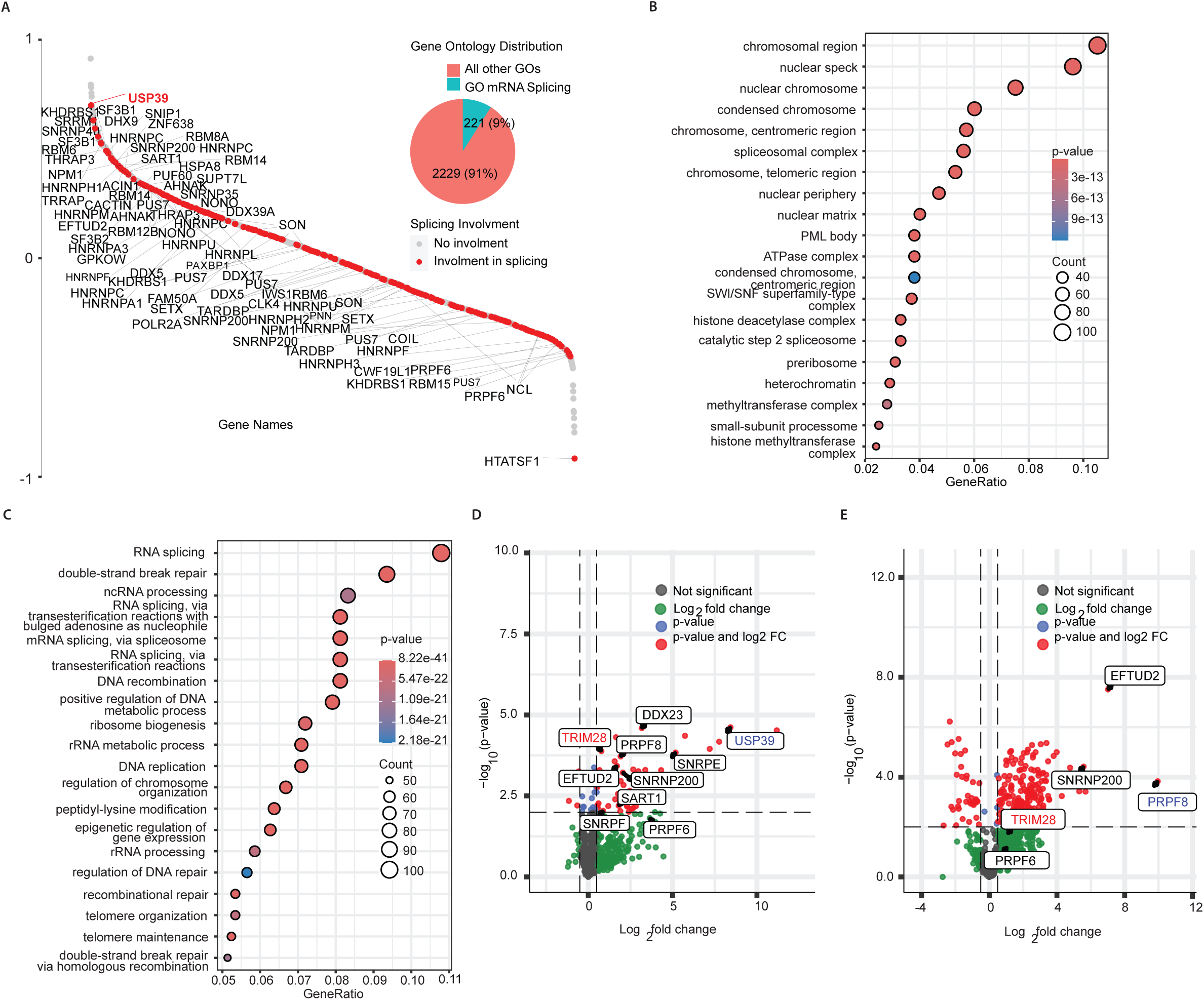
Spliceosomal proteins are heavily SUMOylated and interact with TRIM28. (A) Mass spectrometry-based interactome analysis of SUMO proteins reveals an enrichment of SUMOylated splicing factors. (B) Gene Ontology (GO) enrichment analysis of cellular components among SUMOylated proteins highlights spliceosome-related compartments as top-enriched terms. (C) GO enrichment analysis for biological processes indicates RNA splicing as the most significantly enriched term among SUMOylated proteins. (D) Volcano plot of mass spectrometry data for the interactome of USP39, a spliceosomal protein, shows TRIM28 among the top significantly enriched interactors. (E) Volcano plot of mass spectrometry data for the interactome of PRPF8, a core spliceosomal component, also identifies TRIM28 as a highly enriched interactor.

In order to identify key regulators, that are important for SUMOylation processes of substrate spliceosomal proteins, we decided to use USP39 for further analyses. Interestingly, in the analysis of previously published interactome from mass spectrometry analysis^39^ we identified TRIM28 as a prominent interaction partner of USP39 (Figure 1D). Moreover, by performing co-IP of endogenous PRPF8 followed by MS analysis we also identified TRIM28 (Figure 1E). Based on these data, we selected TRIM28 for further analyses of spliceosome regulation and alterations of splicing processes.

### TRIM28 mediates the SUMO2-driven SUMOylation of USP39

To analyze the post-translational modifications of the spliceosomal protein marker USP39, we performed a series of Immunoprecipitation assays (IPs) and Western blots, which showed an enriched signal for SUMO2-modified USP39 proteins (Figure 2A). Furthermore, we tested SUMOylation specificity by enriching and immunoprecipitating SUMO2 and SUMO3 proteins in cells overexpressing GFP-tagged wildtype USP39 or a USP39 mutant in three lysine residues in then-terminal non-structured part of USP39 that were previously identified as SUMOylated^57^(lysines mutated to arginines, K29R, K51R, K73R, Figure 2B). In our mass spectrometry SUMO-interactome, we identified USP39 as highly SUMOylated splicing-related protein (Figure 1A). For SUMO2 and SUMO3, we detected strong co-precipitation with the wildtype USP39 with no significantly detectable precipitation for the mutant USP39 construct (Figure 2C). Notably, only wildtype USP39 displayed higher-molecular-weight bands detected with anti-His antibodies for detection of His-SUMOylated proteins, indicating enhanced SUMOylation, whereas no such bands appeared with the lysine-to-arginine USP39 mutant.

**Figure 2.**
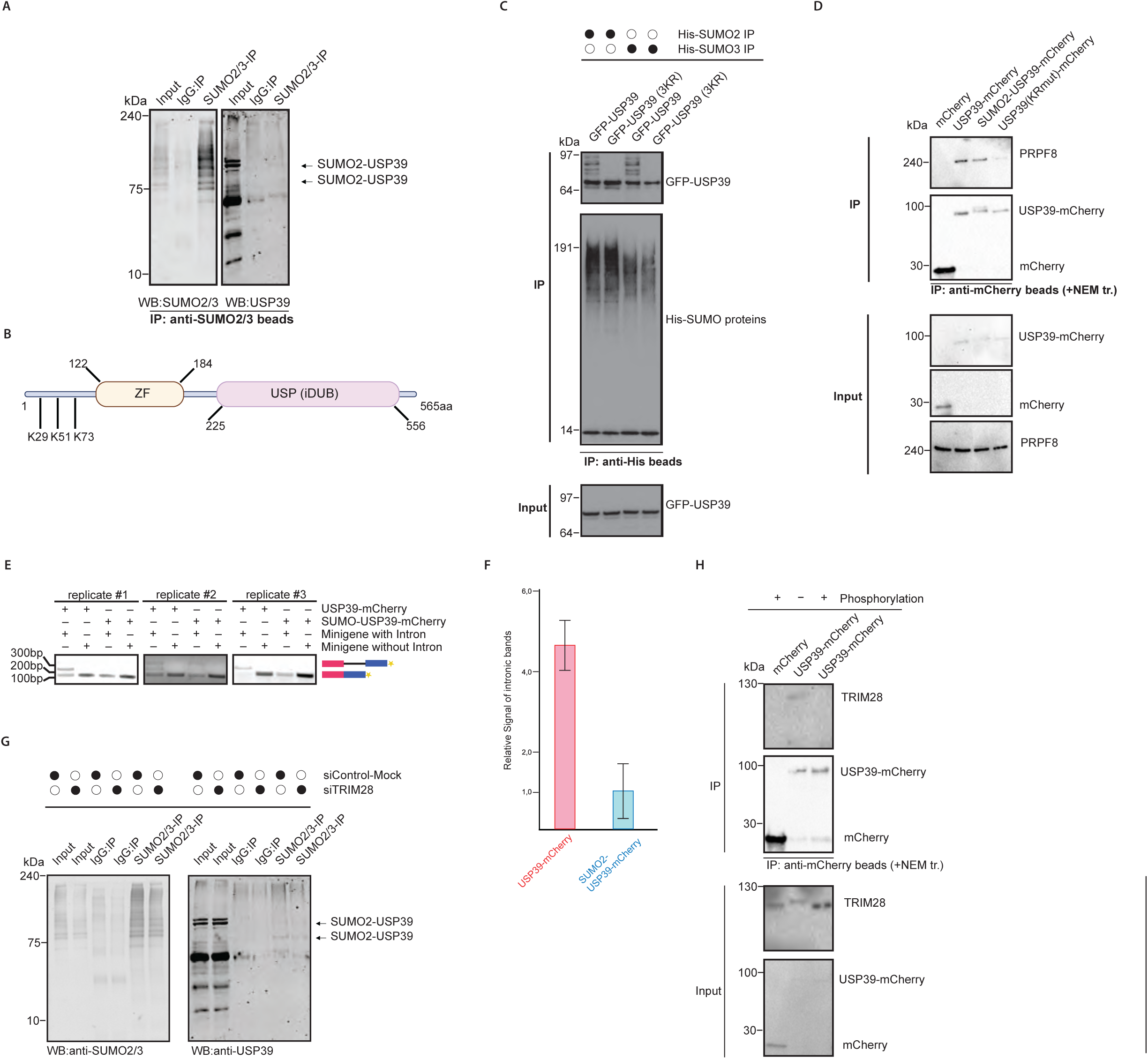
TRIM28 mediates SUMO2-driven SUMOylation of USP39. (A) Western blot analysis of SUMO2/3 immunoprecipitations reveals strong enrichment of SUMOylated USP39. (B) Schematic representation of the USP39 protein domain structure highlighting identified SUMOylated lysine residues. (C) Western blot shows loss of USP39 SUMOylation upon mutation of three key lysines to arginine (K→R), indicating these sites are major SUMOylation targets. (D) Western blot analysis of immunoprecipitated USP39 reveals reduced binding to spliceosomal proteins in the K→R SUMOylation-deficient mutant. (E) Agarose gel electrophoresis of RT-PCR products from a minigene splicing assay shows reduced intronic signal upon expression of SUMOylated USP39, suggesting enhanced intron removal. (F) Quantification of spliced and unspliced PCR bands from the minigene assay shown in (E). (G) Western blot demonstrates decreased SUMOylation of USP39 upon TRIM28 knockdown, implicating TRIM28 as a mediator of E3 SUMO ligase activity. (H) Western blot shows that TRIM28 binds USP39 preferentially in its non-phosphorylated state, suggesting phosphorylation status influences their interaction.

To examine the effect of SUMOylation on USP39’s interaction with spliceosomal proteins, we performed co-IP assays with mCherry-tagged SUMO-fused, wildtype and mutant USP39 (lysines K29R, K51R, K73R from 1-122aa (N-terminal unstructured part of USP39) were mutated to arginines). Once we preserved maximal levels of SUMOylated forms of USP39 by using NEM (N-ethylmaleimide) treatment in the lysis, the USP39-mCherry mutant showed very low binding to the tri-snRNP spliceosomal marker proteins PRPF8, whereas mCherry-tagged SUMO-fused and SUMOylated mCherry-tagged USP39, bound effectively to these proteins, indicating that SUMOylation enhances interaction inside of tri-snRNP complex between USP39 and following spliceosomal proteins. (Figure 2D).

We also performed a minigene splicing experiment using PCR to evaluate intron retention. To this end, we transfected the HEK293T cells with either USP39-mCherry or SUMO2-USP39-mCherry plasmids together with minigene plasmids with p120 exon 5 and 6^58^ with or without the intervening intronic region. In cells expressing mCherry-tagged SUMO-fused-USP39 protein, intronic retention was less detected in semi-quantitative PCR reaction compared to cells expressing USP39-mCherry (Figure 2E,F).

In order to check influence of TRIM28 on the SUMOylation levels of USP39, we performed an IP for detection of SUMOylated proteins in control vs. knockdown of TRIM28, as showed in Figure 2G. While we detected USP39 as SUMOylated protein in control experiments, in case of knockdown of TRIM28, we observed less SUMOylated USP39 in immunoprecipitation experiments, respectively (Figure 2G).

We performed Immunoprecipitation experiments of overexpressed control mCherry and respective USP39-mCherry plasmid in HEK293T cells in the presence or absence of a phosphatase inhibitor (pSTOP) to assess binding of endogenous TRIM28 to USP39 under different phosphorylation states. We observed that TRIM28 interacts with USP39 only in its non-phosphorylated state, while the phosphorylated form of TRIM28 showed no significant binding to spliceosomal protein USP39 as shown in Figure 2H.

### TRIM28 binds to USP39 in phosphorylation-dependent manner revealed by Mass Spectrometry and Venus BiFC Immunofluorescence assays

To further investigate the role of phosphorylation of TRIM28 and interaction with USP39 in phosphorylation-dependent manner in live cells, we cloned a Venus BiFC (bimolecular fluorescence complementation of Venus protein separated in two complementary parts where one part was cloned with TRIM28 and the remaining part cloned with USP39 gene) constructs to monitor binding events of TRIM28 to USP39 in nuclei of live cells (Scheme of Venus constructs illustrated in Figure 3A). In Figure 3B, immunofluorescence images show Venus complexes resulting from TRIM28-USP39 interactions, with localization exclusively in the nucleus, as demonstrated by colocalization with nuclear DAPI staining. The complexes also colocalized with PRPF8, a spliceosomal tri-snRNP marker, in the nucleus. Additionally, IP assays demonstrated that the TRIM28-USP39 Venus complex binds to the spliceosome, interacting with U5 spliceosomal proteins PRPF8 and EFTUD2 (Figure 3C). Interestingly, when the cells were treated with the phosphatase inhibitor orthovanadate to maintain phosphorylation, we observed high levels of phosphorylated TRIM28 with no Venus complex formation, indicating that elevated TRIM28 phosphorylation disrupts TRIM28-USP39 binding (Figure 3D).

**Figure 3.**
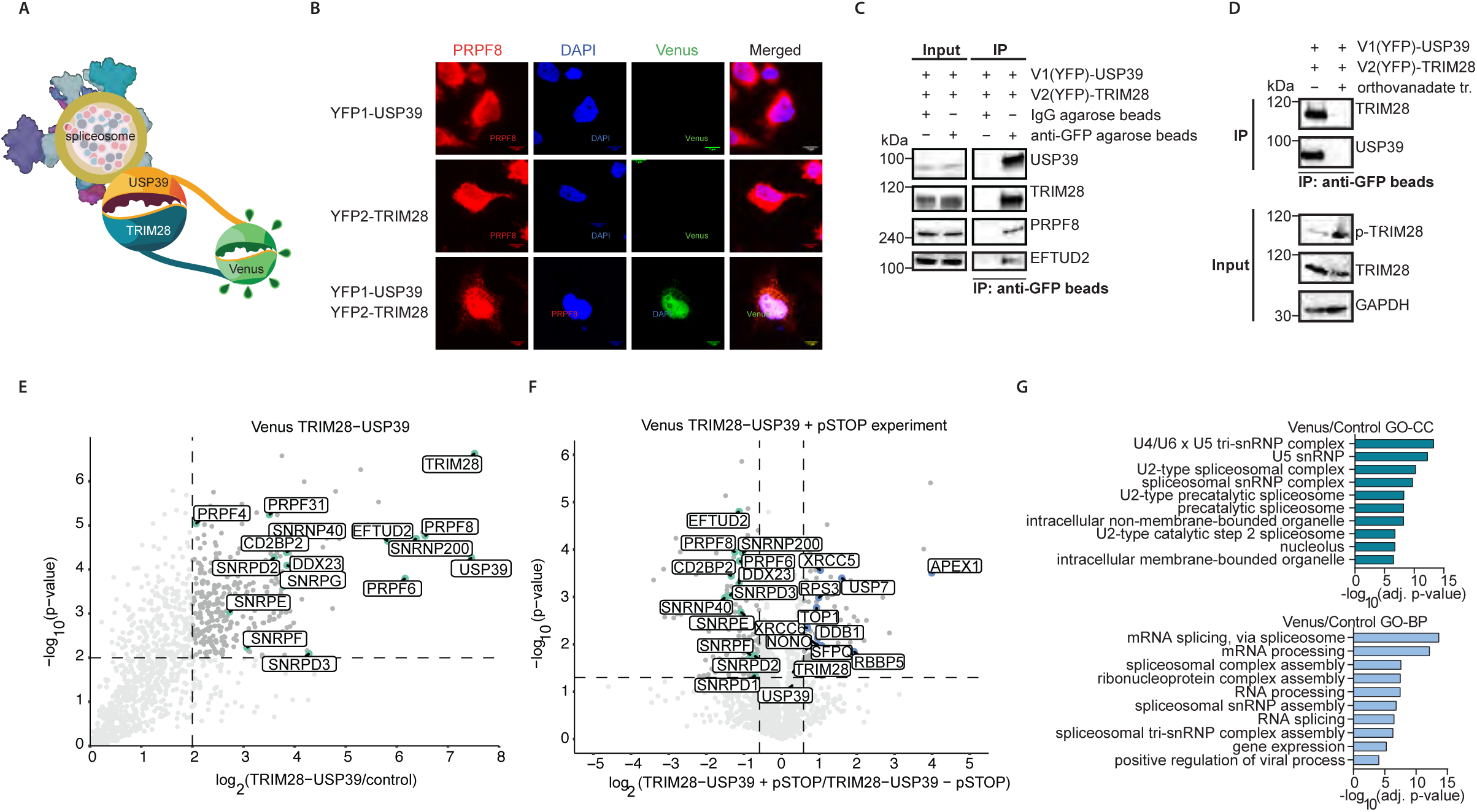
TRIM28 binds to spliceosomal proteins in a phosphorylation-dependent manner. (A) Schematic of Venus-tagged constructs used to investigate TRIM28–USP39 complex formation and interactions with spliceosomal proteins. (B) Representative immunofluorescence images showing subcellular localization of Venus-tagged constructs. (C) Western blot analysis demonstrating that the TRIM28–USP39 Venus complex associates with additional spliceosomal proteins, confirming spliceosome binding. (D) Western blot shows that immunoprecipitation of the Venus complex is detectable only in the absence of phosphorylated TRIM28, indicating a phosphorylation-dependent interaction. (E) Mass spectrometry-based interactome of the Venus complex reveals enrichment of spliceosomal components. (F) Mass spectrometry analysis of the Venus complex upon treatment with phosphatase inhibitors (pSTOP) shows decreased spliceosomal binding and increased association with DNA damage response proteins. (G) Gene Ontology enrichment analysis highlights spliceosomal complexes as the top-enriched cellular components (upper panel) and RNA splicing as the top biological process (lower panel) associated with Venus complex interactors.

To further characterize these interactions, we performed mass spectrometry-based interactome analysis on non-phosphorylated and phosphorylated Venus complexes in cells using the phosphatase inhibitor (pSTOP). As shown in Figure 3E, the non-phosphorylated complex was highly enriched with other U5 spliceosomal proteins, confirming the role of TRIM28-USP39 in spliceosome association. Interactome analysis of the pSTOP-treated samples indicated an enrichment of DNA damage response proteins in the phosphorylated state, with spliceosomal proteins downregulated in this condition (Figure 3F). GO analysis of the proteins identified in the interactome of non-phosphorylated state (Figure 3G) revealed a strong enrichment of splicing protein complexes as the primary cellular components in the non-phosphorylated state (Figure 3G, upper image). Biological process GO analysis further highlighted RNA splicing and mRNA processing as highly enriched pathways among the upregulated proteins identified in the interactome of non-phosphorylated state (shown in Figure 3G, lower image).

### TRIM28 Influences on mRNA splicing

In order to understand the influence of TRIM28 on mRNA splicing, we performed knockdown experiments and RNA sequencing followed by alternative splicing analysis using MAJIQ^59^ tools. We observed alternative splicing events that significantly changed upon TRIM28 knockdown, with intron retention being the most abundant class (Figure 4A,B). Comparison with the alternative splicing events changed upon USP39 knockdown^39^ (Figure 4C) revealed only a limited overlap between the two conditions. Moreover, among the shared alternative splicing events, only about half were changing in the same direction (Figure 4D-F). Together, these observations suggest that TRIM28 impacts mRNA splicing by broader means than just through the USP39 SUMOylation.

**Figure 4.**
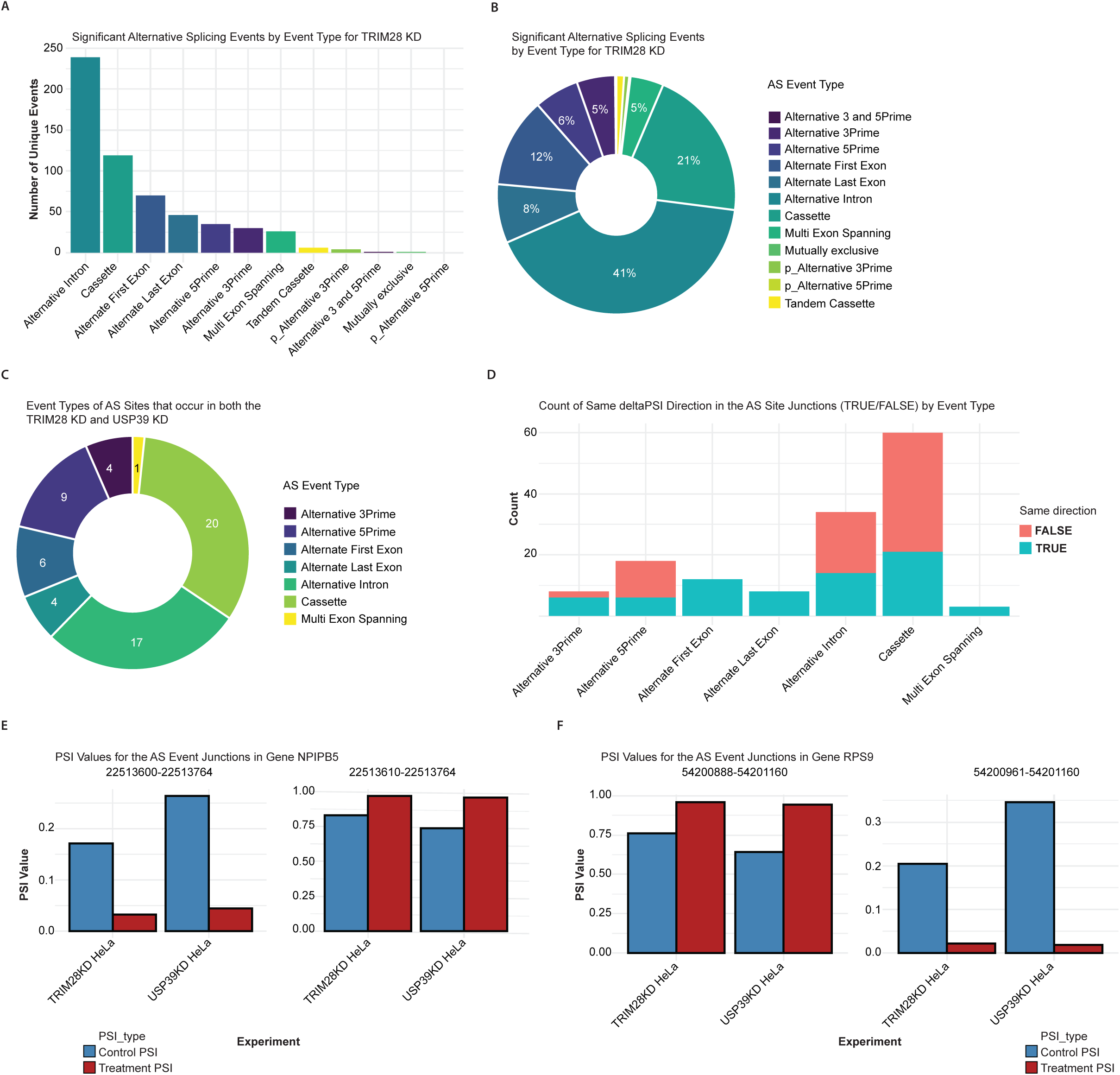
TRIM28 influences mRNA splicing. (A) Bar chart showing the number of significant alternative splicing (AS) events categorized by unique event types (e.g., exon skipping, intron retention) following TRIM28 knockdown. (B) Pie chart representing the proportion of each unique AS event type among all significant events for TRIM28 knockdown dataset. (C) Pie chart showing the distribution of AS event types that are shared between TRIM28 and USP39 knockdown datasets. (D) Bar chart displaying the number of AS events with consistent ΔPSI direction (percent spliced in) across TRIM28 and USP39 knockdowns, categorized by event type. (E) PSI values for the AS junctions of NPIPB5 in control (blue) and knockdown (red) conditions for TRIM28 and USP39 datasets, highlighting similar splicing changes upon depletion of either factor. (F) PSI values for the AS junctions of RPS9 in control (blue) and knockdown (red) conditions for TRIM28 and USP39 datasets, highlighting similar splicing changes upon depletion of either factor.

## Discussion

In this study, we identified a previously unrecognized role for TRIM28 in the regulation of pre-mRNA splicing, mediated through its control of the SUMOylation of the spliceosomal protein USP39. Among spliceosomal components, USP39 emerged as one of the most highly SUMOylated proteins, exhibiting the strongest enrichment in SUMO modification. Notably, SUMOylated USP39 displayed significantly enhanced binding to other spliceosomal proteins, suggesting that this modification promotes its integration or stability within the spliceosome. While previous studies have shown that SUMOylation is widely involved in the DNA-damage response, oxidative stress^60^ as well as in transcriptional regulation^61^, its direct involvement in pre-mRNA has remained incompletely understood. Our findings extend this knowledge by demonstrating that SUMOylation of USP39 facilitates its function, potentially by stabilizing protein-protein interactions within the U4/U6.U5 tri-snRNP complex.

Importantly, we also uncover an additional layer of regulation involving a phosphorylation-dependent interaction between TRIM28 and USP39. TRIM28 preferentially associates with USP39 under non-phosphorylated conditions, without interfering with the assembly of core spliceosomal components. Conversely, upon phosphorylation, TRIM28 no longer binds USP39 and instead redirects its activity toward DNA damage response pathways, consistent with its established role in genome integrity maintenance These findings reveal a dynamic regulatory mechanism whereby TRIM28 toggles between two distinct cellular functions, alternative splicing and DNA repair, based on its phosphorylation status. Given that previous studies have shown TRIM28 is phosphorylated at serine 473 and serine 824 in response to DNA damage, it is plausible that this modification acts as a molecular switch governing TRIM28’s functional partitioning between RNA processing and genome surveillance.

Our minigene splicing assay provided direct evidence that SUMOylation of USP39 influences the efficiency of intron removal. Specifically, we observed reduced intron retention upon expression of SUMOylated USP39, indicating that this post-translational modification may modulate spliceosome activity. Mechanistically, SUMOylation could facilitate recruitment of additional splicing factors or promote conformational changes within the spliceosome that accelerate splicing catalysis, potentially by stabilizing key protein-protein interactions.

Furthermore, our interactome studies highlight the context-dependent role of TRIM28 in spliceosome function. Under non-phosphorylated conditions, TRIM28 associates with USP39 and other core U5 snRNP proteins without interfering with the canonical spliceosomal protein interactions. In contrast, when phosphorylated, TRIM28 loses its affinity for spliceosomal proteins and instead preferentially binds to DNA damage response factors. This behavior aligns with previous reports showing that various genotoxic stresses trigger specific phosphorylation events in key proteins such as p53^62,63^. This dual functionality suggests that TRIM28 acts as a molecular switch, integrating signals from distinct cellular pathways to modulate splicing efficiency.

Given that alternative splicing is a central mechanism for regulating gene expression, our findings offer insights into how post-translational modifications can influence splicing fidelity. Notably, TRIM28 dysregulation has been implicated in multiple cancers and neurodegenerative disorders, conditions frequently characterized by aberrant splicing patterns. It will be important to determine whether TRIM28-mediated SUMOylation of USP39 is altered in disease context and whether this interface between splicing and DNA damage pathways could be exploited for therapeutic purposes.

In summary, our study identifies TRIM28 as new regulator of spliceosomal function through mediating SUMOylation of USP39. We propose a new conceptual framework in which post-translational modifications, such as SUMOylation, play a pivotal role in orchestrating gene expression. Future investigations should address the broader impact of TRIM28-driven SUMOylation on splicing outcomes across physiological and pathological settings.

## Acknowledgments

We acknowledge Antonia Hofmann and Karin Siegmund for technical support at Institute of Biochemistry II (IBC2; Goethe University, Frankfurt). We thank all members of the Quantitative Proteomics Unit at IBC2 (Goethe University, Frankfurt), in particular, Georg Tascher, Martin Adrian-Allgood for technical help and measurements, Kristina Wagner for preparing LC columns and David Krause for help in (bio)informatics. The authors also thank Isabel Hahl for all additional help in creating plasmid constructs and technical help in western blot experiments.

We acknowledge funding from the Deutsche Forschungsgemeinschaft (DFG, German Research Foundation) – SFB1361.

## Author contribution

Conceptualization, V.M., C.P.-G. and I.D.; Methodology, V.M., C.P.-G., T.M., P.H., N.B., K.Z., M.M.M., A.C.O.V., S.M. and I.D.; Investigation, V.M., C.P.-G, T.M., N.B., J.S. and P.H.; Writing V.M. and I.D. with contribution of other co-authors; Funding Acquisition, I.D.; Resources, I.D., K.-Z., S.M., M.M.M. and W.G.; Supervision, I.D.

---

**Supplementary Figure S1. Analysis of the SUMO proteome, related to Figure 2**.

(A) Table with highlighted Top 45 highest SUMOylated proteins in human proteome with normalized log2 intensities

**Supplementary Figure S2.**
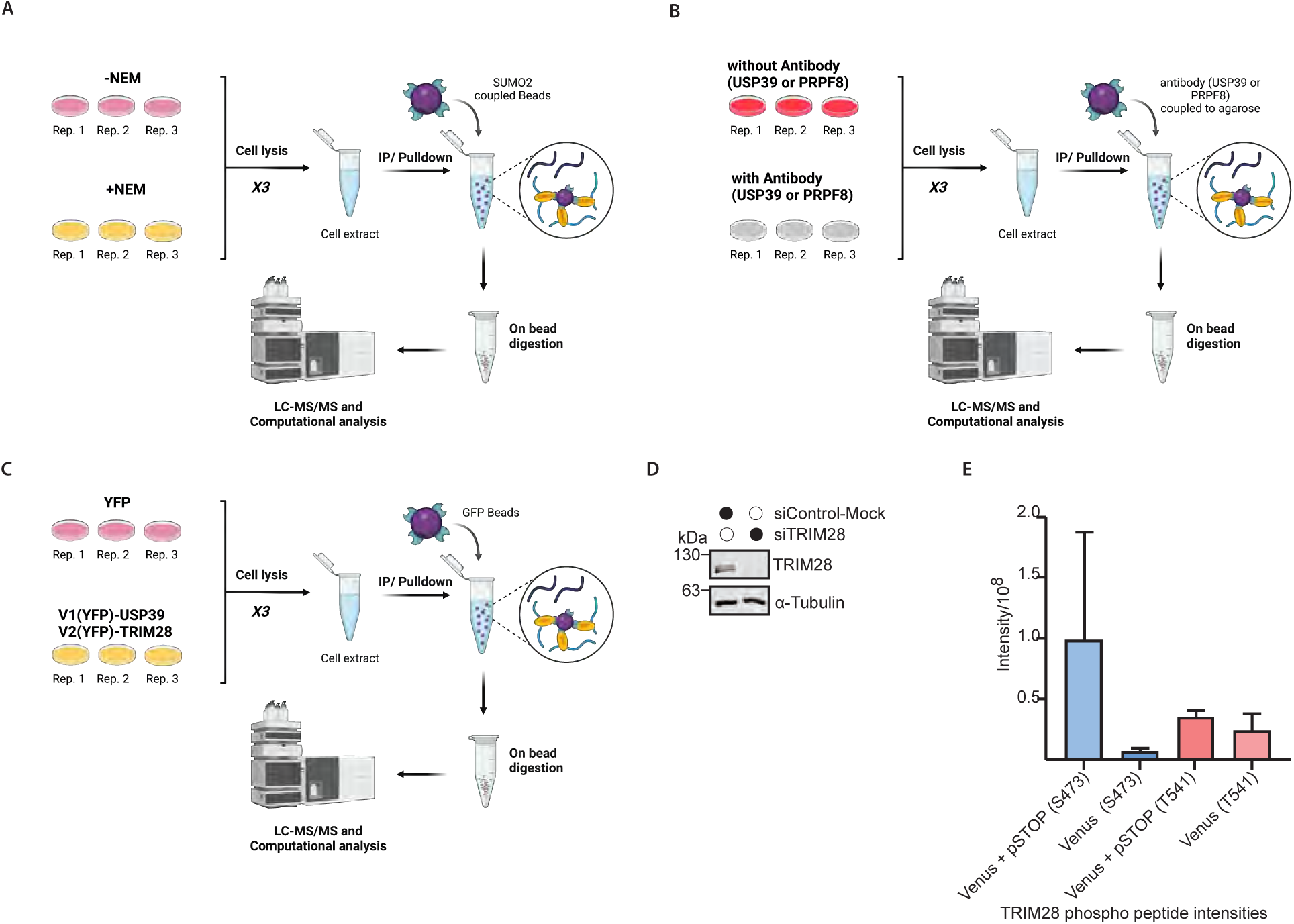
Schematic representation of Mass spectrometry SUMO proteome experiments and USP39 & PRPF8 and Venus interactome experiments with analysis of relevant phospho sites in Venus complex upon pSTOP treatment, related to Figures 2. (A) Schematic representation of mass spectrometry SUMO proteome analysis (B) Schematic representation of mass spectrometry USP39 & PRPF8 interactome analysis (C) Schematic representation of mass spectrometry Venus BiFC interactome analysis. (D) Western blot showing TRIM28 knockdown in siTRIM28-treated cells compared to siControl-Mock (Control experiment for Figure 2G, α-Tubulin serves as a loading control). (E) TRIM28 phospho peptide intensities for major phospho sites detected in Venus BiFC TRIM28-USP39 interactome analysis.

## STAR METHODS

### LEAD CONTACT AND MATERIAL AVAILABILITY

Further information and requests for reagents should be directed to and will be fulfilled by the Lead Contact Ivan Dikic. All reagents generated in this study are available from the Lead Contact following the completion of a material transfer agreement (MTA).

### DATA AVAILABILITY STATEMENT

The datasets presented in this study can be found in online repositories. The names of the repository/repositories and accession number(s) can be found below: tba

### EXPERIMENTAL MODEL AND SUBJECT DETAILS

#### Mammalian cells

HeLa and HEK 293T cells were obtained from the American Type Culture Collection (ATCC). Cells were cultivated in Dulbecco’s modified Eagle’s medium (DMEM) supplemented with 10% fetal bovine serum with Penicillin Streptomycin Solution (Gibco, according to manufacture protocol). All cells were cultured at 37 °C in a humid 5% carbon dioxide atmosphere. All cells used in this study were tested negative for mycoplasma contamination.

### METHOD DETAILS

#### Cell lines culture and transfection

HEK293T, HeLa cells were purchased from ATCC. Cells were cultured in high glucose Dulbecco’s Modified Eagles Medium (DMEM) supplemented with 10% fetal bovine serum (FBS), 100 U/mL penicillin and 100 mg/mL streptomycin at 37 °C, 5% CO2 in a humidified incubator. Plasmids were introduced into HEK 293T cells using Lipofectamine 2000 as the transfection reagent according to manufacturer’s protocol. Lipofectamine RNAiMAX was used for the introduction of siRNA into HEK 293T and HeLa cells according to manufacturer’s protocol. Pre-designed siRNAs were purchased from Merck.

#### Western blotting and immunoprecipitation

Cell lysates or immunoprecipitated proteins were mixed with SDS sample buffer (1% SDS in 50 mM Tris-HCl, pH 8), heated at 95 °C for 5 min, centrifuged, and separated by Tris-Glycine SDS-PAGE, and transferred to PVDF membrane (Millipore) at cold room. Blots were blocked with 5% nonfat milk for 1 h at room temperature and incubated with primary antibodies overnight at cold room or 2 h at room temperature and washed with TBST (0.1% Tween 20 in TBS) three times. The blots were further incubated with secondary antibodies for 1 h at room temperature and washed 3 times with TBST. The blots were incubated with ECL reagents (Advansta), and chemiluminescence was acquired with the Bio-Rad ChemiDoc system. For immunoprecipitation, HEK293T cells expressing GFP-tagged, GFP-similarly created proteins (Venus BiFC complex^64^) or endogenous cells were lysed with mild immunoprecipitation buffer containing 150 mM NaCl, 50 mM Tris-HCl, pH 7.5, 0.5% NP40, 1 mM PMSF, protease inhibitor cocktail (Sigma Aldrich), mixed with 10 μL GFP-trap, RFP-trap, or antibody conjugated agarose beads, and incubated for 4 h in cold room with end-to-end rotation. Beads were washed 3 times in IP buffer containing 500 mM NaCl. Proteins were eluted by resuspending with 2X SDS sample buffer followed by boiling for 5 min at 95 °C. Samples were then submitted to western blotting analysis.

#### Immunofluorescence

HeLa cells were seeded on a coverslip in 12-well plates and cultured in CO2 incubator. Next day cells were transfected with plasmids for Venus BiFC constructs. Briefly, cells were washed once with PBS, pH 7.4, and fixed with 4% paraformaldehyde (PFA) in PBS for 10 min at room temperature. Cells were washed again with PBS 2 times, then permeabilized with 0.1% Triton-X100 in PBS for 10 min, and blocked with blocking buffer containing 0.1% saponin and 2% BSA in PBS for 1 h at room temperature. Cells were stained with antibodies diluted in blocking buffer overnight at 4 °C and washed with PBS three times next day. Cells were further incubated with DAPI in PBS, followed with further 3 times washing with PBS. Confocal imaging was performed using the Zeiss LSM780 microscope system. Images were analyzed with Fiji software.

#### Chemical treatment of the cells

All of the chemical compounds used in treatments were prepared in 1000X stock to the working concentration, and in order to not reach more than 0,1% of solvent volume percentage. Control mock cells were always treated with respective solvent in equal amounts.

#### Mass spectrometry

##### Mass spectrometry sample preparation (co-immunoprecipitation)

Immunoprecipitation experiments were performed as described in section “Western blotting and Immunoprecipitation” with additional steps of 3 times washing the beads with PBS after washing steps with 500 mM sodium chloride buffer in order to remove all salts from the beads before elution. Proteins were eluted from the beads by adding elution buffer (2% sodium deoxycholate, 1mM TCEP, 4 mM CAA, 50 mM ammonium bicarbonate pH 8.5) and boiling at 95°C for 10 min. The supernatant was removed before briefly adding dilution buffer (50 mM ammonium bicarbonate pH8.5) to the beads and combining the supernatant with the first elution. 500 ng of LysC/Trypsin each was added o/n at 37°C. Isopropanol containing 1% TFA was added to stop the digestion prior to purifying the peptides using SDB-RPS tips. Peptides were dried and resuspended in 2% ACN, 0.1% TFA before LC-MS/MS analysis.

##### Mass spectrometry data acquisition (co-immunoprecipitation)

Samples were analysed on a Q Exactive HF coupled to an easy nLC 1200 (ThermoFisher Scientific) using a 35 cm long, 75µm ID fused-silica column packed in house with 1.9 µm C18 particles (Reprosil pur, Dr. Maisch), and kept at 50°C using an integrated column oven (Sonation). Peptides were eluted by a non-linear gradient from 4-28% acetonitrile over 75 minutes and directly sprayed into the mass-spectrometer equipped with a nanoFlex ion source (ThermoFisher Scientific). Full scan MS spectra (350-1650 m/z) were acquired in Profile mode at a resolution of 60,000 at m/z 200, a maximum injection time of 20 ms and an AGC target value of 3*10^6^ charges. Up to 10 most intense peptides per full scan were isolated using a 1.4 Th window and fragmented using higher energy collisional dissociation (normalised collision energy of 27). MS/MS spectra were acquired in centroid mode with a resolution of 30,000, a maximum injection time of 110 ms and an AGC target value of 1*10^5^. Single charged ions, ions with a charge state above 5 and ions with unassigned charge states were not considered for fragmentation and dynamic exclusion was set to 20s.

#### Mass spectrometry data analysis

MS raw data were analyzed using MaxQuant^65^ (v1.6.17.0) (PMID: 21254760, PMID: 27809316). Label-free co-IP experiments were analyzed with the in-build label-free quantification algorithm MaxLFQ using default parameters (PMID: 24942700). Acquired spectra were searched against the human reference proteome (Taxonomy ID 10090) downloaded from UniProt (14-02-2023; 20,206 entries, one sequence per gene) and a collection of common contaminants using the Andromeda search engine integrated in MaxQuant (PMID: 21254760, PMID: 27809316). Identifications were filtered to obtain false discovery rates (FDR) below 1% for both peptide spectrum matches (PSM; minimum length of 7 amino acids) and proteins using a target-decoy strategy (PMID: 17327847). For all searches, carbamidomethylated cysteine was set as a fixed modification and oxidation of methionine, N-terminal protein acetylation, STY phosphorylation, and NEM on cysteines as variable modifications with allowing up to 5 modifications per peptide. Trypsin/P was set as specific proteolytic enzyme with allowing up to two missed cleavages.

#### siRNA-mediated TRIM28 knockdown

siRNA-mediated knockdown of TRIM28 in HeLa cells was performed using a predesigned TRIM28 siRNAs (Merck). One day prior to transfection, HeLa cells were plated at 20% confluency in 10cm plates. The next day, lipofection was performed using RNAiMAX (Thermo Fisher Scientific) according to manufacturer’s instructions. Briefly, 20 μM siRNAs was mixed with OptiMEM (Thermo Fisher Scientific) in one reaction and lipofectamine RNAiMAX was mixed with OptiMEM in a second reaction. Both reactions were incubated at room temperature (RT) for 5 min and mixed by flicking. Meanwhile, culture media on HeLa cells was changed to 10 ml DMEM + 10% FBS lacking antibiotics. Contents of both reactions were mixed and further incubated for 20 min at RT. Finally, 1mL of lipofection mix were added drop-wise onto HeLa cells. As a control, scrambled-sequence of siTRIM28 was used as described for the TRIM28 siRNAs. The transfected cells were incubated for 48 h at 37°C and 5% CO2 and subsequently analyzed in downstream assays.

#### Splicing assay

HEK293T cells were transfected with p120 plasmids (with only exons for control plasmids & plasmids with artificial exons and introns) and SUMO/WT USP39-mCherry tagged plasmids accordingly to transfection protocol from manufacturer (ThermoFisher, Lipofectamine 2000). After two days, RNA was extracted (Macherey-Nagel) and reversibly transcribed to cDNA (ThermoFisher) according to manufactures protocols. cDNAs were diluted and used as template in PCR experiments (Initial denaturation 98 °C for 30 seconds followed by 30 cycles (98 °C for 20 seconds, 67 °C for 30 seconds and 72 °C for 30 seconds) and final extension at 72 °C for 10 minutes. Samples were run on agarose gel (100V, 1h, 2% agarose) and imaged with UV light.

## Key resources table

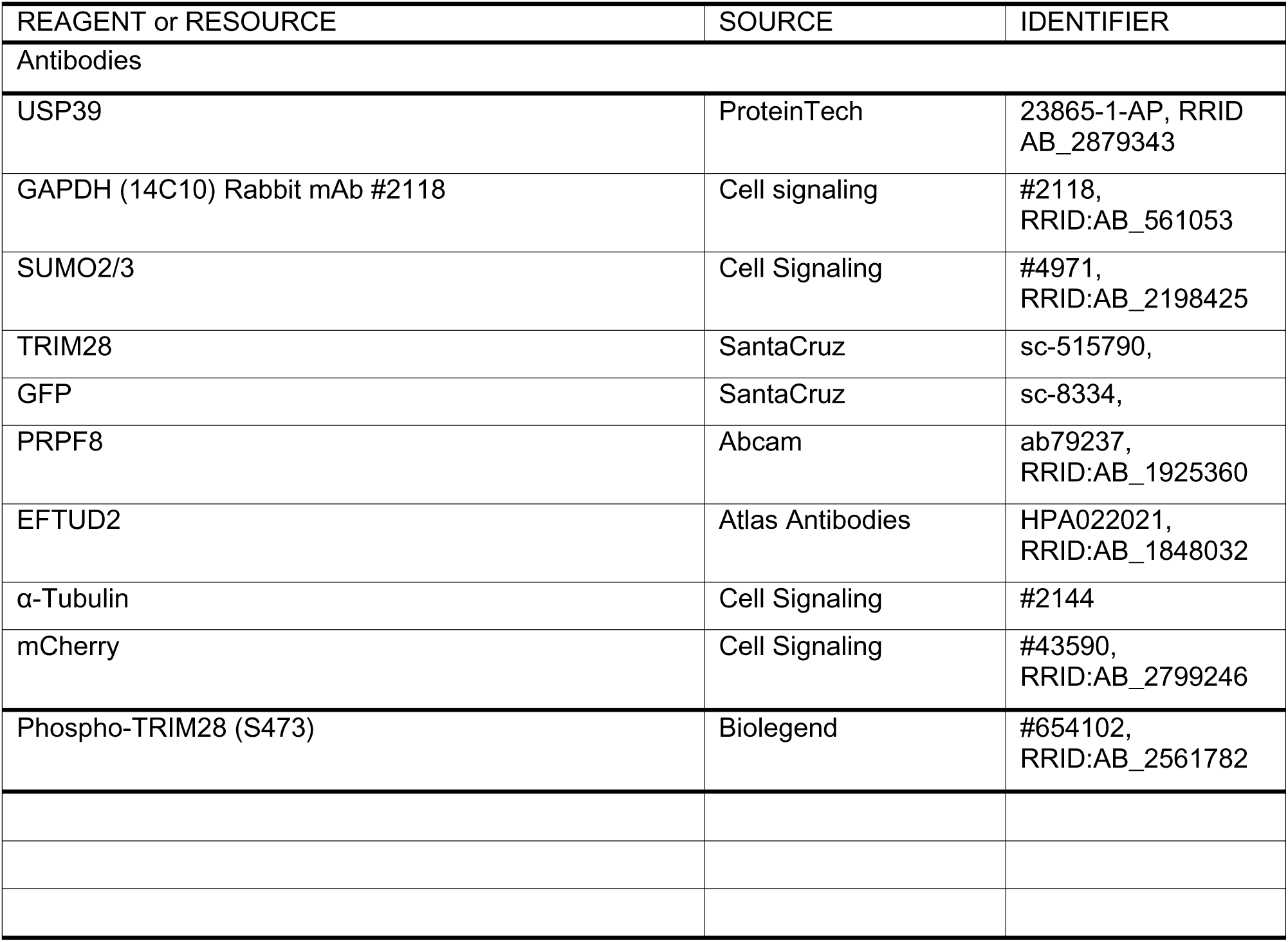
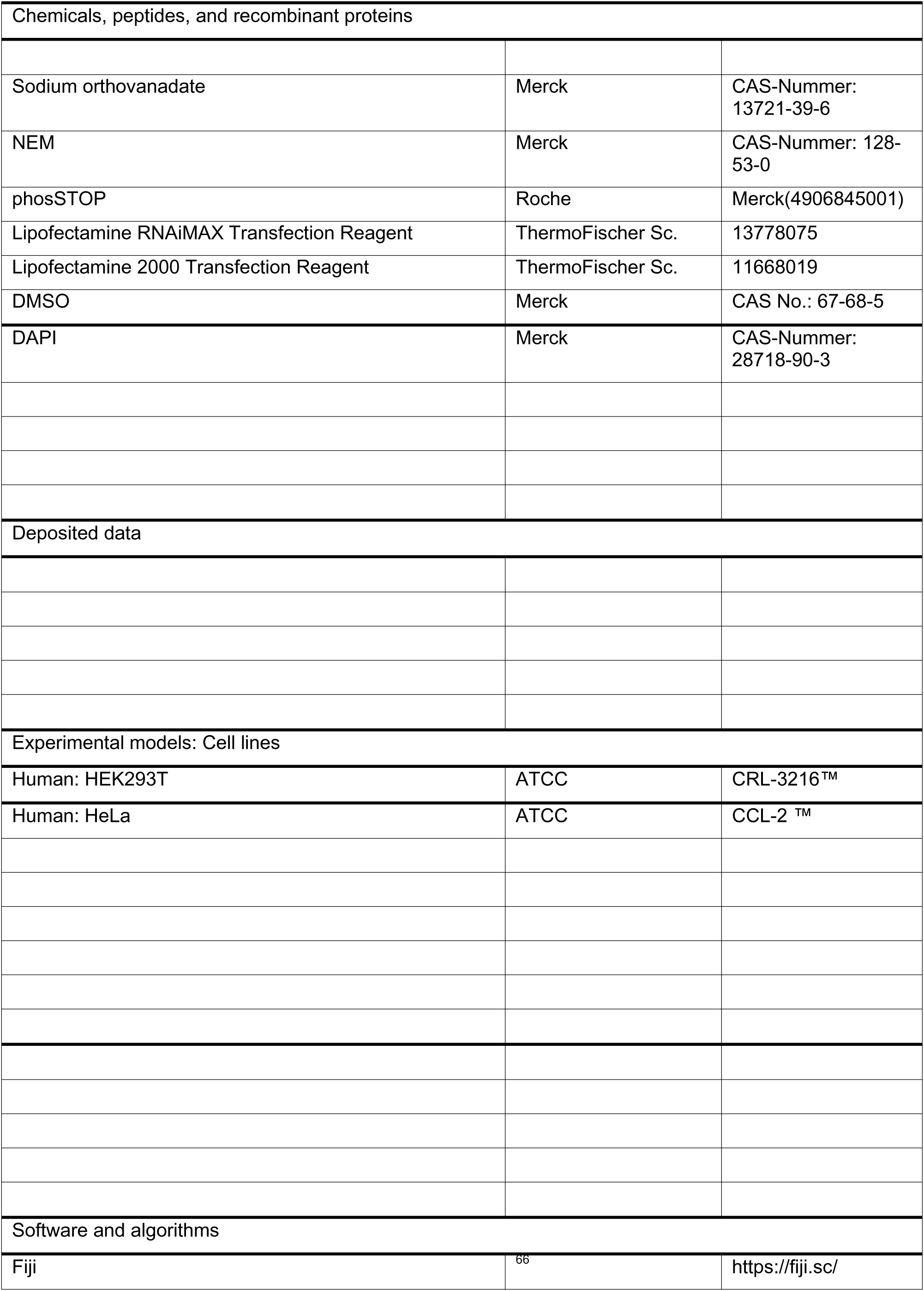
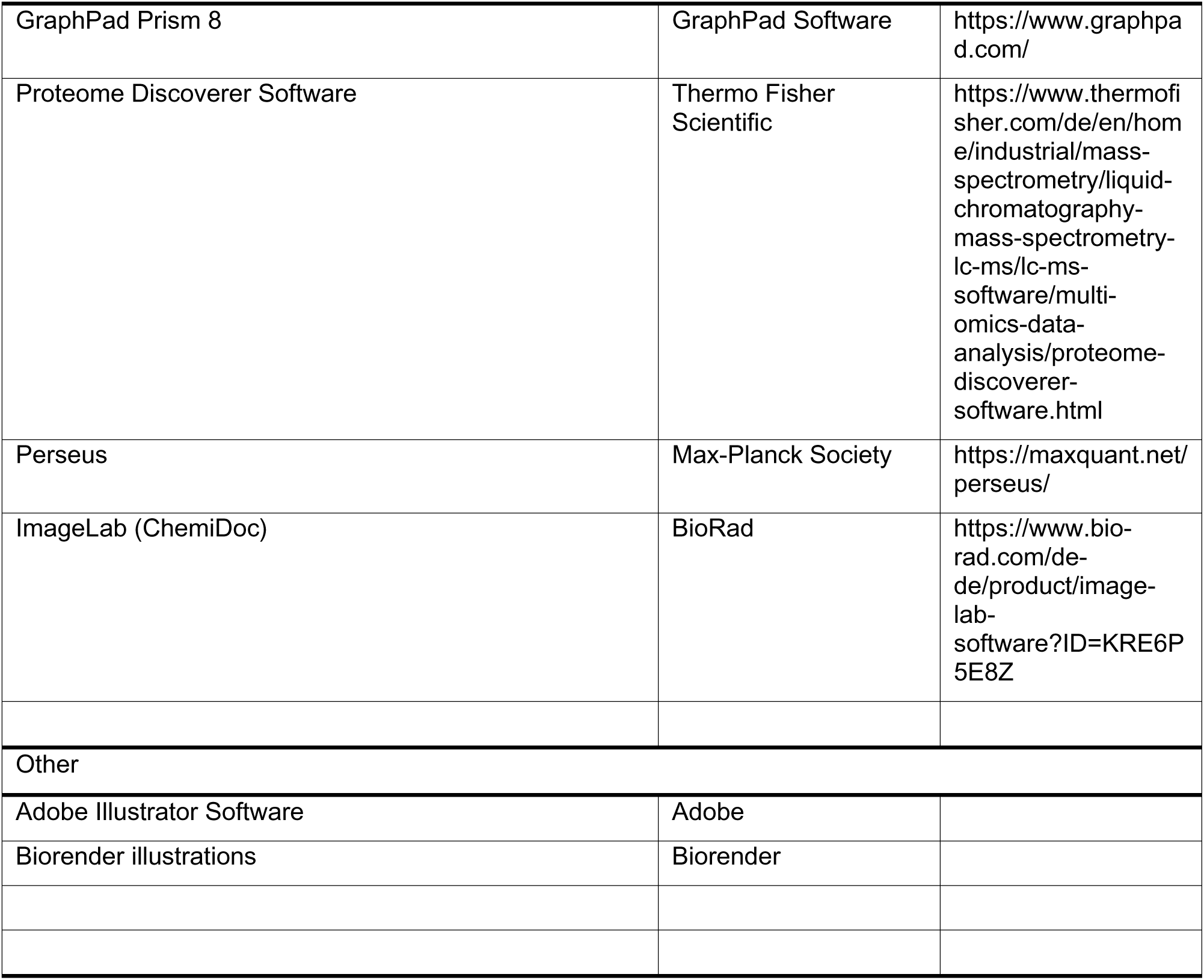

---

**Supplementary Table S1.**
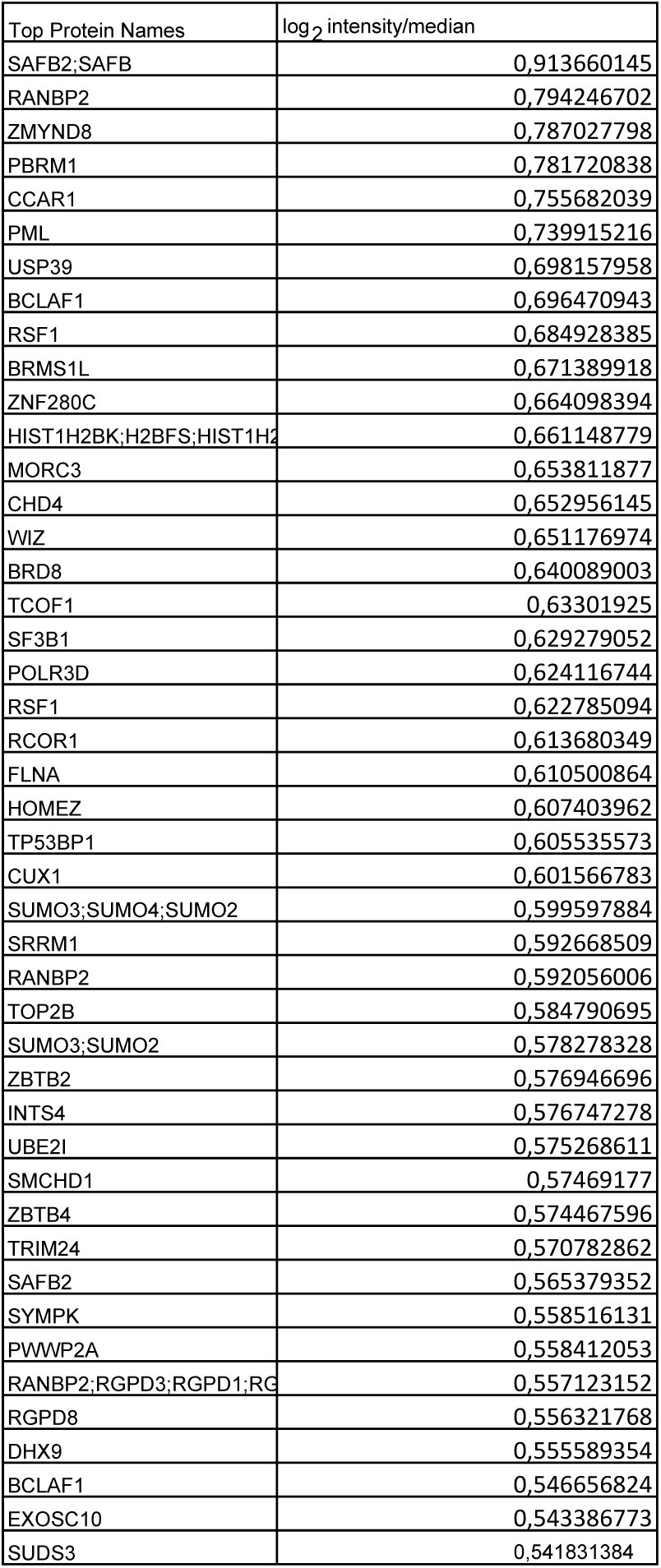
Top 45 highest SUMOylated proteins in human proteome with normalised log2 intensities.

| Top Protein Names | log<sub>2</sub> intensity/median |
| --- | --- |
| SAFB2;SAFB | 0,913660145 |
| RANBP2 | 0,794246702 |
| ZMYND8 | 0,787027798 |
| PBRM1 | 0,781720838 |
| CCAR1 | 0,755682039 |
| PML | 0,739915216 |
| USP39 | 0,698157958 |
| BCLAF1 | 0,696470943 |
| RSF1 | 0,684928385 |
| BRMS1L | 0,671389918 |
| ZNF280C | 0,664098394 |
| HIST1H2BK;H2BFS;HIST1H2A | 0,661148779 |
| MORC3 | 0,653811877 |
| CHD4 | 0,652956145 |
| WIZ | 0,651176974 |
| BRD8 | 0,640089003 |
| TCOF1 | 0,63301925 |
| SF3B1 | 0,629279052 |
| POLR3D | 0,624116744 |
| RSF1 | 0,622785094 |
| RCOR1 | 0,613680349 |
| FLNA | 0,610500864 |
| HOMEZ | 0,607403962 |
| TP53BP1 | 0,605535573 |
| CUX1 | 0,601566783 |
| SUMO3;SUMO4;SUMO2 | 0,599597884 |
| SRRM1 | 0,592668509 |
| RANBP2 | 0,592056006 |
| TOP2B | 0,584790695 |
| SUMO3;SUMO2 | 0,578278328 |
| ZBTB2 | 0,576946696 |
| INTS4 | 0,576747278 |
| UBE2I | 0,575268611 |
| SMCHD1 | 0,57469177 |
| ZBTB4 | 0,574467596 |
| TRIM24 | 0,570782862 |
| SAFB2 | 0,565379352 |
| SYMPK | 0,558516131 |
| PWWP2A | 0,558412053 |
| RANBP2;RGPD3;RGPD1;RGPD2 | 0,557123152 |
| RGPD8 | 0,556321768 |
| DHX9 | 0,555589354 |
| BCLAF1 | 0,546656824 |
| EXOSC10 | 0,543386773 |
| SUDS3 | 0,541831384 |

